# Functional-Hybrid Modeling through automated adaptive symbolic regression for interpretable mathematical expressions

**DOI:** 10.1101/2021.06.09.447664

**Authors:** Harini Narayanan, Mariano Nicolas Cruz Bournazou, Gonzalo Guillén-Gosálbez, Alessandro Butté

**Author notes:** **Corresponding Author:** Alessandro Butté, DataHow AG, Zurich, Switzerland, **E-mail:**.

## Abstract

Mathematical models used for the representation of (bio)-chemical processes can be grouped into two broad paradigms: white-box or mechanistic models, completely based on knowledge or black-box data-driven models based on patterns observed in data. However, in the past two-decade, hybrid modeling that explores the synergy between the two paradigms has emerged as a pragmatic compromise. The data-driven part of these have been largely based on conventional machine learning algorithm (e.g., artificial neural network, support vector regression), which prevents interpretability of the finally learnt model by the domain-experts. In this work we present a novel hybrid modeling framework, the Functional-Hybrid model, that uses the ranked domain-specific functional beliefs together with symbolic regression to develop dynamic models. We demonstrate the successful implementation of these hybrid models for four benchmark systems and a microbial fermentation reactor, all of which are systems of (bio)chemical relevance. We also demonstrate that compared to a similar implementation with the conventional ANN, the performance of Functional-Hybrid model is at least two times better in interpolation and extrapolation. Additionally, the proposed framework can learn the dynamics in 50% lower number of experiments. This improved performance can be attributed to the structure imposed by the functional transformations introduced in the Functional-Hybrid model.

## 1. Introduction

The plethora of mathematical models in science and engineering available can be broadly classified into two paradigms: (i) data-driven, statistical or (Machine Learning (ML)) models, and (ii) first principle based (mechanistic, white box) models. Both approaches have their own advantages and disadvantages as summarized in [1,2] and a choice is made based on the prior understanding about the system and the availability of data. In chemical engineering [3–9] and biotechnology [1,10–16], hybrid modeling is emerging as a pragmatic solution to mathematical modeling, exploring the synergy between the two paradigms. Hybrid models have been very successful in systems that are only partially understood, and the availability of data is limited or/and costly.

Most of the hybrid modeling work pose basic mass or energy balances and some preliminary dependencies and approximate unknown relationships with data-driven models. The most popular data-driven model used in such frameworks is a shallow (typically single layer) artificial neural network [6,8,9,13–16] with some literature also reporting use of subspace identification algorithms [17], non-linear Partial Least Squares [18], Adaptive regression splines [19], Support Vector Machine (for regression) [20,21], Gaussian Processes [22] and Recurrent Neural Network [7].

However, such machine learning algorithms lead to hybrid models that are hard to interpret. In addition, there are also common domain expectation of functional forms that can be accounted for to further introduce expert knowledge in the hybrid modeling framework. For instance, in chemical reaction kinetics, typically power law models are expected or in biochemistry and biochemical network models hill kinetic type equations are predominant. Similarly, in cell culture applications, monod-type or haldane-kinetic equations are expected for metabolites. Incorporating these domain expert considerations in a strategic manner, while still ensuring flexibility, is essential to develop more robust hybrid models while allowing some interpretability from the final hybrid model.

One possible manner to achieve this is through symbolic regression, which attempts to simultaneously identify the model structure and the parameters defining the structure [23]. Symbolic regression has been applied in different fields such as chemical systems [23–25], fluid dynamics [26], and natural science [27,28], for the purpose of data-driven system identification. The symbolic regression problem can be tackled by applying genetic programming over a system of arithmetic operators (+, -, x, /) and functions (identity (), log (), exp (), trigonometric, polynomial) to choose the optimal order of operators represented through the so-called expression trees. An alternative approach to symbolic regression, developed in the recent times, involve the sparse regression method [29–32], which creates a candidate library of all possible functional transformations and their interaction terms and applies a LASSO regression to choose only a subset from the candidate library to best describe the observed data. Successful application of the SINDy based algorithms have been demonstrated for dynamic systems such as in biological networks [29] and to learn partial differential equations as well [32]. Here again simple functional transformations such as *identity (), log (), exp ()*, trigonometric and polynomial with some variant also considering rational functions are used [29].

Still in most engineering applications, the general scope of modelling is to arrive to a mathematical formulation that explains the observations of a system sufficiently well, is as simple as possible and is interpretable by the domain-experts. With this in mind, here we present an approach for symbolic regression that employs functional transformations based on domain-expertise in a systematic procedure. Additionally, we remove the arithmetic operators used in symbolic regression and introduce a multiplicative and additive series (which is a common occurrence in all equations). Finally, instead of using genetic programming we used genetic algorithms (GAs) to select and reject functions. A detailed description of the implementation used in this work can be found in Section 2.2.2. The developed algorithm is first tested on four benchmark systems: (i) Reaction scheme identification of chemical species, (ii) Enzyme kinetics, (iii) Lotka-Volterra problem and (iv) FitzHugh-Nagumo (FHN) problem. Finally, its added value is demonstrated with a microbial fermentation bioreactor. In-silico simulators were used to generate data for all the case studies. First, the ability of the algorithm to accurately represent the data generating equations while providing interpretability to finally converged hybrid model is highlighted for the different studies. Second, the comparison of the Functional-Hybrid model with the equivalent hybrid model framework using artificial neural network (Hybrid-ANN) is presented. Finally, practical implication of the Functional-Hybrid model in terms of number of experiments required and extrapolation capability is highlighted using the microbial bioreactor case study, which are both essential features in engineering applications.

## 2. Materials and Methods

### 2.1. Materials

The proposed algorithm was tested on four benchmark problems and a system of relevance to biotech industry. The conditions used in the simulation of the different cases are described below and tabulated in Table 1, while the detailed equations used in the simulation can be found in the supplementary information.

**Table 1:**
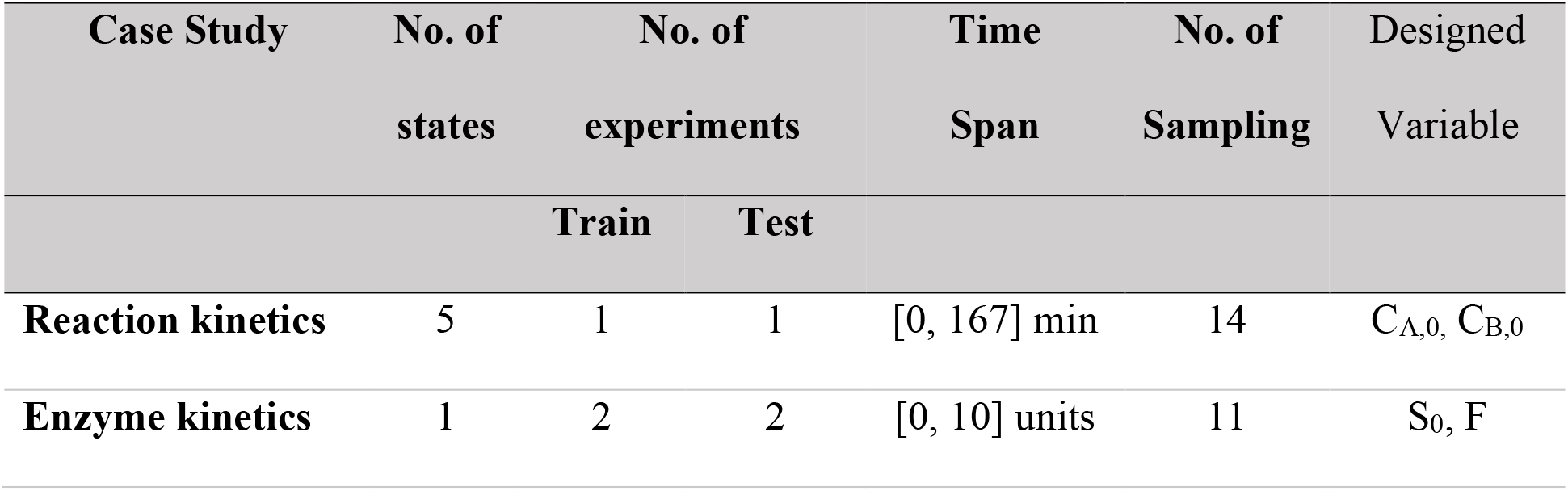
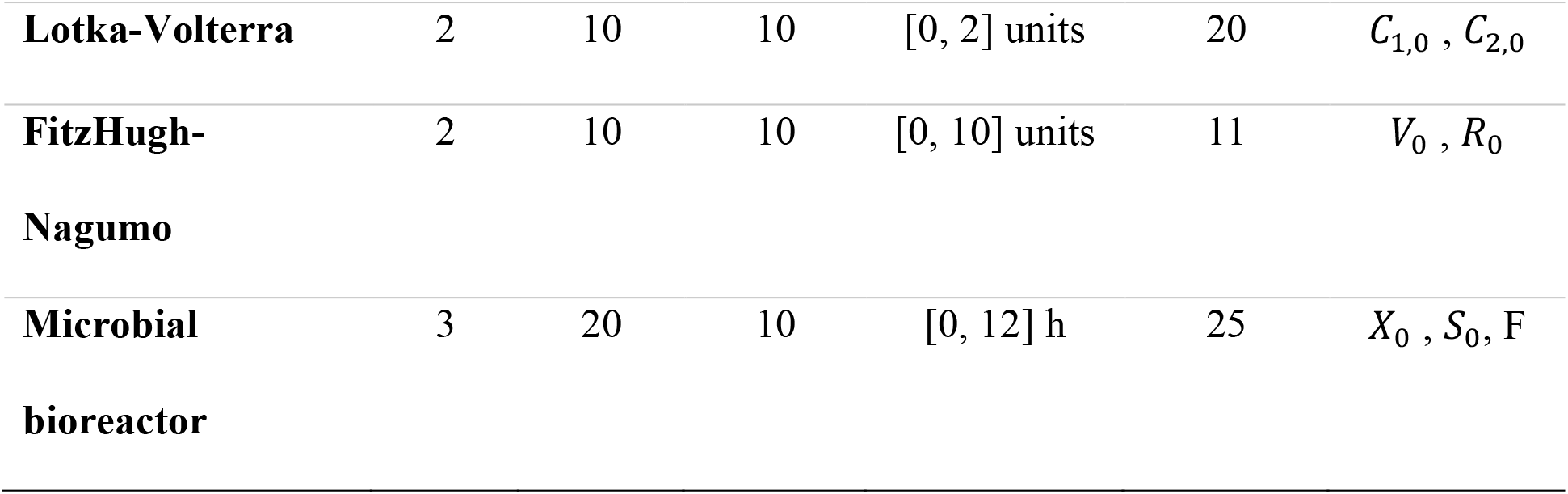
Conditions used in the simulation of the different cases

#### 2.1.1. Chemical reaction kinetics

To study the reaction kinetics of two species A and B, we carried out an in-silico simulation of an experiment at different flowrates and fixed inlet concentration (C_A,0_ = C_B,0_ = 2 mol/L) in a Plug Flow Reactor (PFR) reactor of 500 L of volume. The system consists of 5 species namely, A, B, P, D and S, whose concentration is measured at the outlet. The evolution profile of each of the species, as a function of residence time, is used to predict the reaction kinetics using our algorithm. Data corresponding to 14 different residence times in the interval [0, 167] min are generated for all the 5 species.

#### 2.1.2. Enzyme Kinetics

The Michaelis-Menten model is the most popular equation to describe enzyme kinetics. It captures enzyme binding and unbinding with substrate and subsequent irreversible formation of the product [29]. Time series data of the substrate S (one variable), simulated at two different initial condition (S_0_) and inlet fluxes (F), are used to extract the functional form using our algorithm. A white noise of 5% standard deviation was added to the true profile to generate measurements considering 11 time points in the time interval of [0, 10] units. The established model is then tested on two additional experiments measured at a different initial substrate concentration and inlet flux.

#### 2.1.3. Lotka-Volterra

The third system used in the study is the Lotka-Volterra system originally proposed in [33] and used as benchmark to study algorithms of parameter estimations such as in [34,35]. The problem describes the dynamics of two states (reported in SI), that was used to simulate data. 20 experiments are planned using a Latin hypercube sampling method [36] for the initial condition of the two states (*C*_1,0_, *C*_2,0_). Among the twenty, 10 randomly chosen experiments are used for training and 10 experiments are used for testing the model identified by the algorithm. Time profiles are simulated in the time interval of [0, 2] units and measurements perturbed with 10% gaussian noise at 20 equally space time points are used for modeling

#### 2.1.4. FitzHugh-Nagumo (FHN) model

The FHN models, is a system proposed by FitzHugh [37] and Nagumo [38] for modeling giant squid neurons and often used to test parameter estimation robustness for spiky dynamics [34,35]. The dynamics is represented by a system of two ordinary differential equation (ODE), reported in SI. 20 experiments are planned using a Latin hypercube sampling method for the initial condition of the two states (*V*_0_, *R*_0_) out of which 10 randomly chosen experiments are used for training and 10 experiments are used for testing the model identified by the algorithm. Time profiles are simulated in the time interval of [0, 10] units and measurements perturbed with 10% gaussian noise at 11 equally space time points are used for modeling.

#### 2.1.5. Microbial fermentation bioreactor

A microbial fed-batch bioreactor is simulated using a system of ODE of three variables, namely the biomass, the substrate and the product. The system of equations is adapted from [39] and is reported in the SI. For the base case (discussed in section 3.2), 30 experiments are planned using a Latin hypercube sampling method, varying the feed flowrate (F) and the initial concentration of biomass (*X*_0_) and substrate (*S*_0_). The concentration of feed stream is kept constant at S_f_ = 150 g/L. 20 experiments are used to train the model and 10 experiments are used to test the model. Time profiles are simulated in the interval of [0, 12] h and measurements perturbed with 15% gaussian noise at 25 equally spaced time points are then used for modeling. It is here noted that, for the case study discussed in section 3.3 wherein minimum number of experiments required to develop the hybrid models is evaluated, 50 experiments are simulated using the same procedure stated above. Subsequently. among these, the test set consisting of 10 experiments is randomly chosen and kept fixed. With the remaining 40 runs, training sets of different sizes are prepared with randomly chosen runs.

For the extrapolation case study (discussed in section 3.3), in addition to the 50 experiments, 10 experiments are simulated also using a LHS method but with the initial concentration of all the species being outside the ones used to train the model.

### 2.2. Method

All the simulations and modeling are performed in MATLAB 2019b. For all the models, concentrations normalized to the maximum value of the respective states in the training set are used as inputs. The different models are compared based on the Root Mean Squared Error in prediction (RMSEP), computed with respect to the true values (not the perturbed measurements), of individual states involved in each case study. Additionally, when a single overall metric (aggregating the performance across different runs, time points and state variables) is required to represent model performance, the overall normalized Mean Squared error in prediction (*MSEP*_*norm,max*_) is used. The formula for the *MSEP*_*norm,max*_ calculation is as follows:

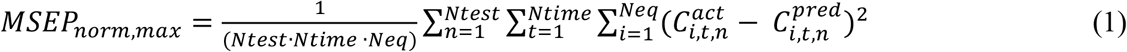

where 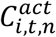 and 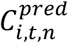 are, respectively, the true and predicted concentration of the *i*-th species at *t*-th time point of the *n*-th run in the test set, normalized to the maximum value of the respective species in the training set. The details about the framework and the implementation of the Functional-Hybrid model and the benchmark hybrid model with ANN (Hybrid-ANN) are presented in the following sections.

#### 2.2.1. Functional-Hybrid model

Functional-Hybrid models are inspired by the common patterns observed in all the formalized equations. In other words, each equation can be decomposed in blocks that are combined additively (addition or subtraction). Subsequently, each block is constituted by certain entities that are multiplied. Finally, these entities can be represented as functional transformations of the system variables including the states representing the system, process conditions (Z) and control variables (W). This is encoded in the Functional-Hybrid model as represented schematically in Figure 1A, and illustrated through an example in Figure 1B. The algorithm takes as input the variables (*X*) and all the possible functional transformations (*f*) to be applied to each variable, together with the ranking of likely functions. Currently, the prior expectations of likely transformations are hardcoded as rankings but in the future, a Bayesian approach with prior probability distributions could be considered to encode these beliefs.

**Figure 1:**
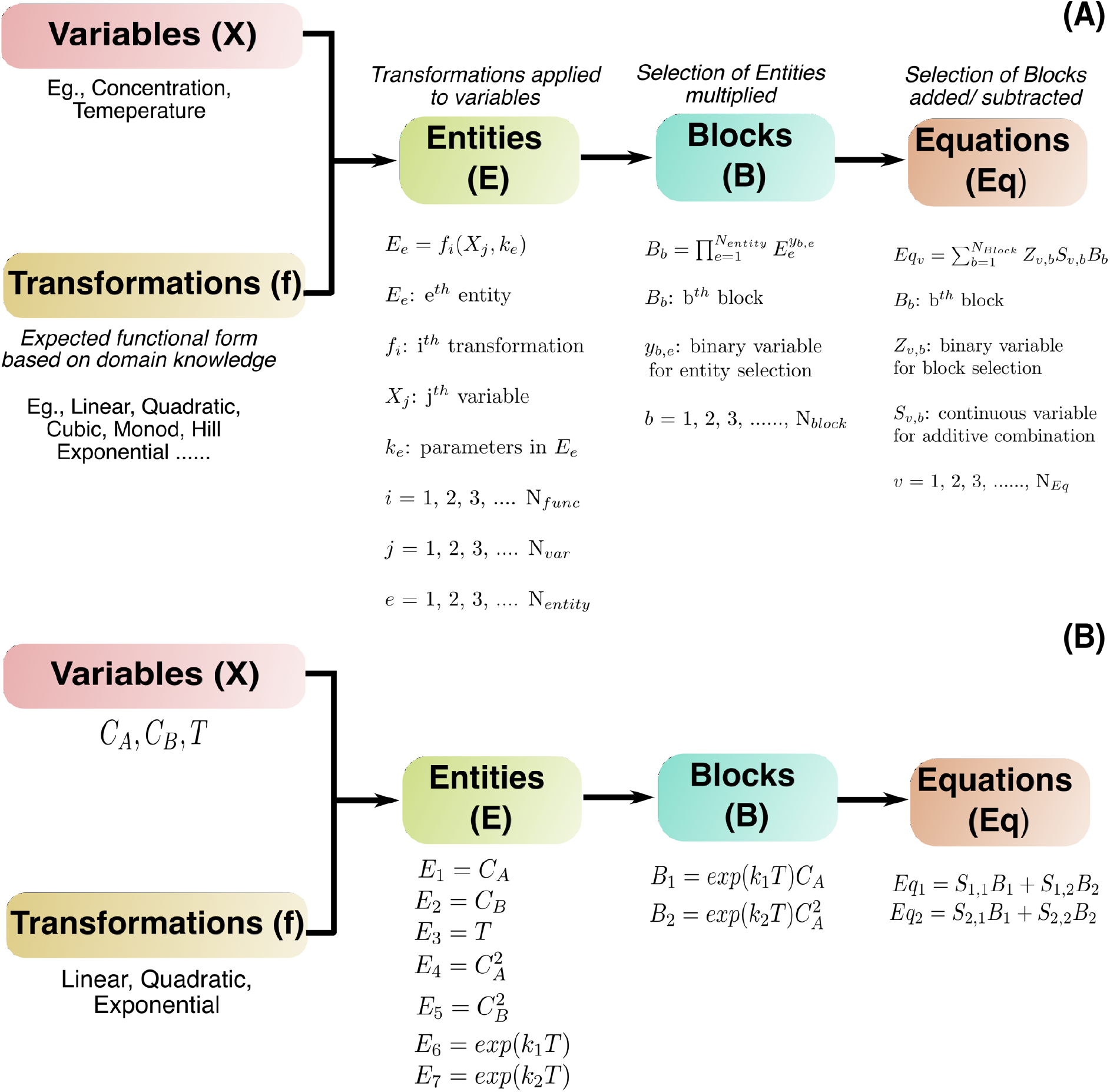
(A) Schematic flowchart of the conceptual workflow and the different components involved in the set-up of the Functional-Hybrid model. (B) Workflow indicated through an example.

The chosen transformations are applied to the respective variables to create the so-called entities, E. These entities then interact in a multiplicative manner to produce the blocks (B) that are then additively combined using coefficients to give the final equations (Eq). These equations could represent the right-hand-side of a dynamic system of ordinary differential equation, as done in all the examples, but can also give rise to a standalone system of algebraic equations.

Overall, the system of equations representing the Functional-Hybrid model are as follows:

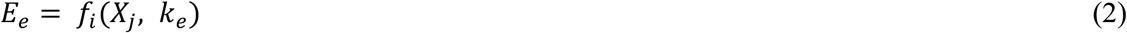

where *E*_*e*_ is the *e-th* entity and *e* = 1, 2, 3, …., *N*_*entity*_. *N*_*entity*_ is defined by the total number of transformations (*f’s*) applied to the different variables (*X*). *k* is the vector of all the parameters used in the different functional transformations and, thus, *k*_*e*_ is a subset of values of the parameters involved in defining entity *E*_*e*_. The entities are combined multiplicatively into blocks as follows:

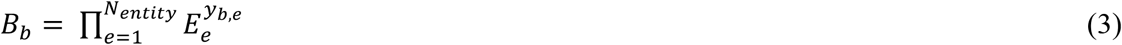

where *B*_*b*_ is the *b-th* block and *b* = 1, 2, 3, …., *N*_*block*_. The number of blocks (*N*_*block*_) is a tuning parameter to be optimized. *y*_*b,e*_ is a binary variable {0,1}, to decide if a certain entity *E*_*e*_ is present in a given block *B*_*b*_. Y is thus a *N*_*block*_ x *N*_*entity*_ matrix of binary variables for all block-entity combination. The blocks are combined additively into equations as follows:

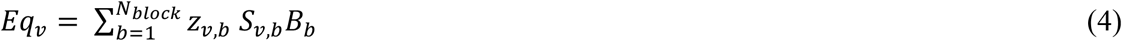

where *Eq*_*v*_ is the *v-th* equation and *v* = 1, 2, 3, …., *N*_*eq*_. The number of equation (*N*_*eq*_) is dictated by the number of states of the system. *Z*_*v,b*_ is a binary variable {0,1} that models if a certain block *B*_*b*_ is present in a given equation *Eq*_*v*_. Subsequently, *S*_*v,b*_ is the continuous variable used in the additive combination of the different blocks, which is zero if *z*_*v,b*_ is zero or is a real value. Z is thus a *N*_*eq*_ x *N*_*block*_ matrix of binary variables for all equation-block combination and S is the corresponding matrix of real-valued coefficient values.

The algorithmic implementation of the Functional-Hybrid model is realized as indicated in the flowchart shown in Figure 2A. The algorithm takes as input the variables (X), functional transformations (f), their corresponding ranking (R) and the options for the tunable parameter *N*_*block*_ (NB). Thereby, the ranking of the functional transformations taken as input from the user is used to set-up a stepwise incorporation. First, the top ranked transformations for each variable are used to define the entities and the first option for *N*_*block*_ is used. Optimization is performed and performance is evaluated based on *MSEP*_*norm,max*_. If the selected settings succeed in meeting the threshold (defined based on the process noise), the algorithm is terminated. If not, optimization is performed using the next option for the tuning parameter, *N*_*block*_. If the performance threshold is not met using the any of the options for *N*_*block*_, the next transformations in the ranking are performed in addition to the already existing ones and the optimization process is repeated. The process terminates when the errors shown by the model reach the pre-determined threshold.

**Figure 2:**
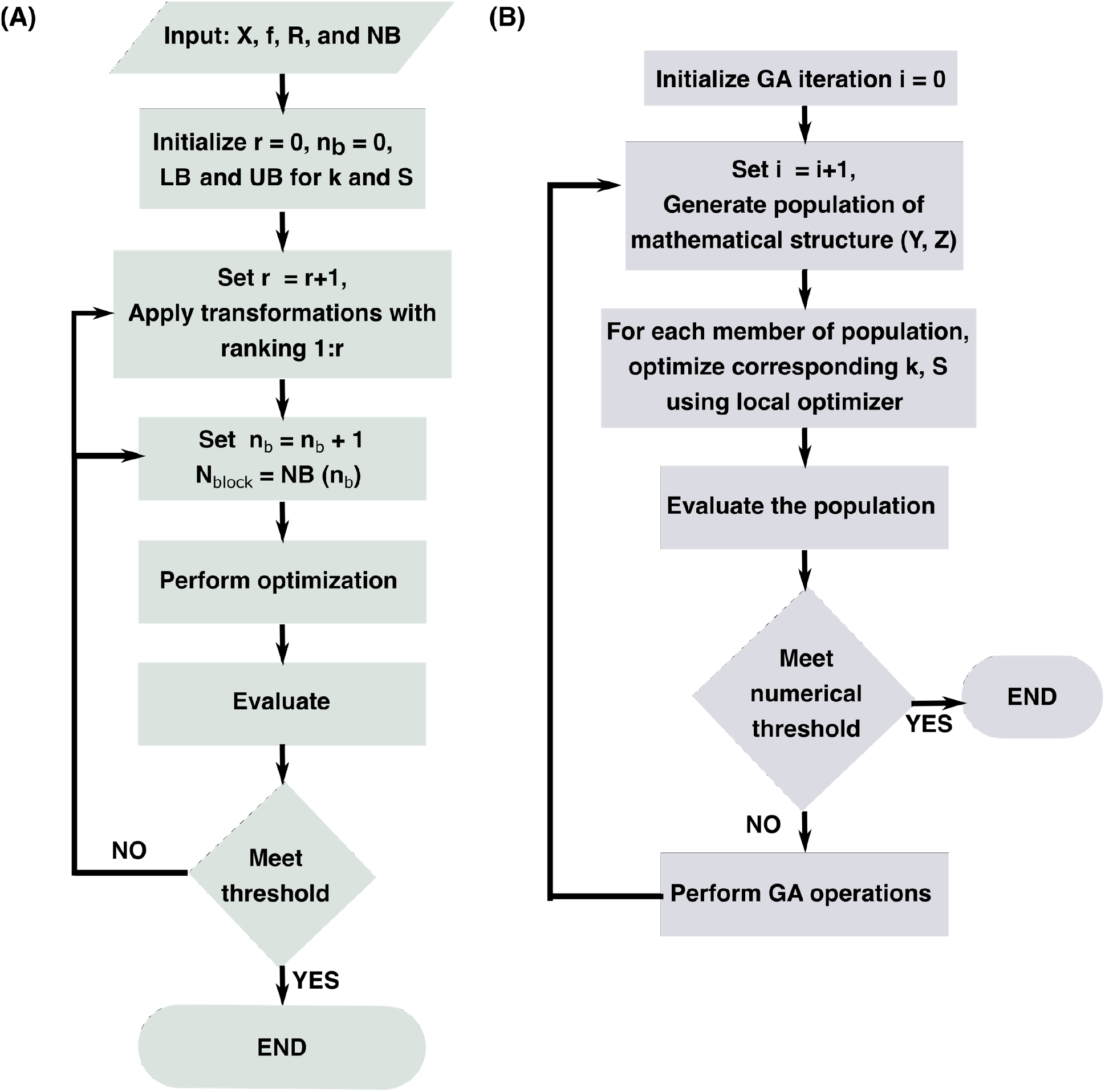
(A) Flowchart representation of the algorithmic workflow of the Functional-Hybrid model. (B) Flowchart representation of the optimization procedure used in the Functional-Hybrid model.

In particular the optimization of the Functional-Hybrid model is implemented by decoupling the discrete optimization for the mathematical structure, dictated by *Y* and *Z*, from the continuous optimization for the real-valued parameters defining the structure, *k* and S. Figure 2B presents a flowchart representation of the sequential optimization procedure used. The outer loop consists of a genetic algorithm (GA), based on the algorithm developed by Deep et al [40] for integer and mixed integer optimization, implemented within the in-built function *ga()*. The GA proposes a population of mathematical structures in every iteration. The continuous optimizer in the inner loop, based on a non-linear programming optimizer (NLP), solves a parameter estimation problem to define the optimal parameters corresponding to each mathematical structure proposed by the GA. The interior-point method implemented in *fmincon()* [41] is used as the NLP optimizer. The inner continuous optimizer in turn calls the numerical integrator based on the Numeric Differentiation Formula (NDF) with *ode15s()* [42], in every iteration to simulate the states and compute the mean squared error (MSE) with respect to the measured experimental values, which serves as the objective function for the NLP. On the other hand, the GA uses as objective the MSE of each proposed mathematical structure with its optimal continuous parameters *(k,S)* determined by the continuous optimizer, and optimizes for the structure definition by tuning the *Y* and *Z*.

#### 2.2.2. Hybrid-ANN model

To illustrate advantages of using the Functional-Hybrid model in practical (or industrial) applications, a comparison is made with the equivalent hybrid model using an artificial neural network, referred to as Hybrid-ANN, for the microbial fermentation bioreactor case study. The equations for the Hybrid-ANN model can be represented as follows:

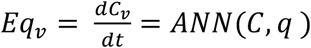

where *C*_*v*_ is the concentration of the *v-th* species that is X, S and P, C is the vector of concentration of all the species and θ is the weight matrix of the ANN. In this specific case study dealing with a microbial fermentation bioreactor, no specific process conditions are varied in the simulation. However, for other cases, a vector of process conditions (e.g., DO, pH, Temperature) is additionally provided as input to the ANN. A multi-output feedforward ANN is used in this work with a *tanh* activation function. Integration and optimization are performed simultaneously to optimize the ANN weights such that the difference between measured and model predicted concentration is minimized. *ode15s ()* and *fminunc ()* is used for integration and optimization, respectively [1].

## 3. Results and Discussion

### 3.1. Algorithm Validation studies

Firstly, the performance of the proposed algorithm was studied on four simple benchmark systems: (i) identification of the reaction kinetics from dynamic measurements of the reactant and product, (ii) identification of the enzyme kinetics, (iii) Lotka-Volterra system and (iv) FHN system. The functional-Hybrid model is trained as described in section 2.2.1 on the training set defined in section 2.1 for the respective case study. The trained (or optimized) model is then applied on the respective test data to compute the RMSEP and the *MSEP*_*norm,max*_. The supplementary information (S.I. Table 1 through 5) provides the details of the different ranked transformations, the converging iteration following the step-wise addition approach, and the number of optimal *N*_*blocks*_ used in each of the four case-studies.

For an exemplary run in the test set, Figure 3 presents the comparison of the dynamic evolution of the concentration measured (crosses) and predicted (dashed lines) in the different case studies. The RMSEP observed on each state for the respective case-study is tabulated in Table 2. Both the dynamic profile and the RMSEP indicate that the Functional-Hybrid model is capable of accurately capturing the true trends and is minimally influenced by the measurement noise. Additionally, for these cases, the system of ODE equations learnt by the Functional-Hybrid model was very close to the original system of ODE used to simulate the data. Table 2 shows the final equations deduced by the Functional-Hybrid model for the four benchmark case studies and the original in-silico model equations used to simulate data. It is noteworthy that the continuous parameters (k and S) of the converged equations do not exactly match the respective in-silico model since the Functional-Hybrid model, unlike the original model, is trained on normalized concentrations.

**Table 2:**
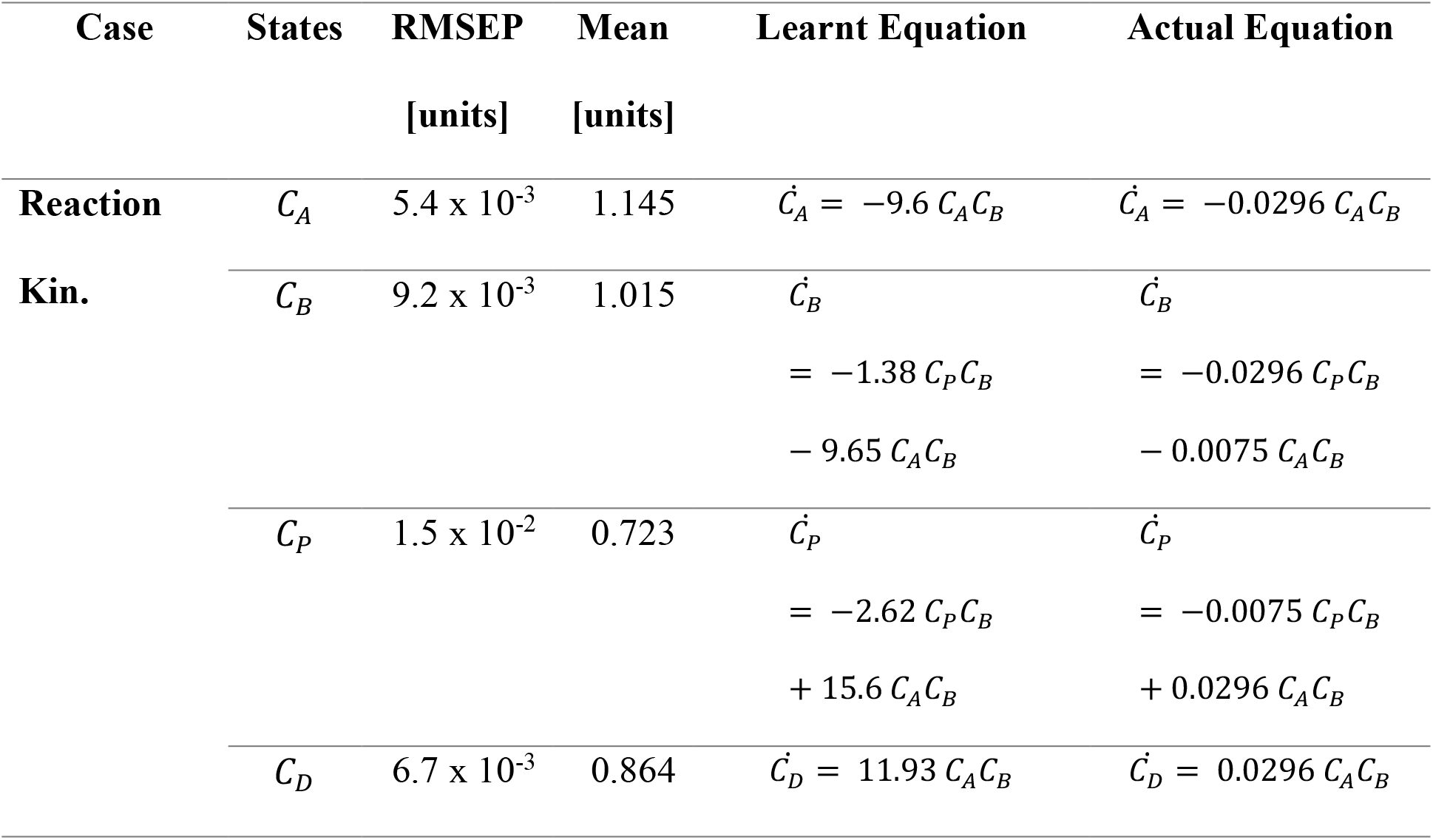
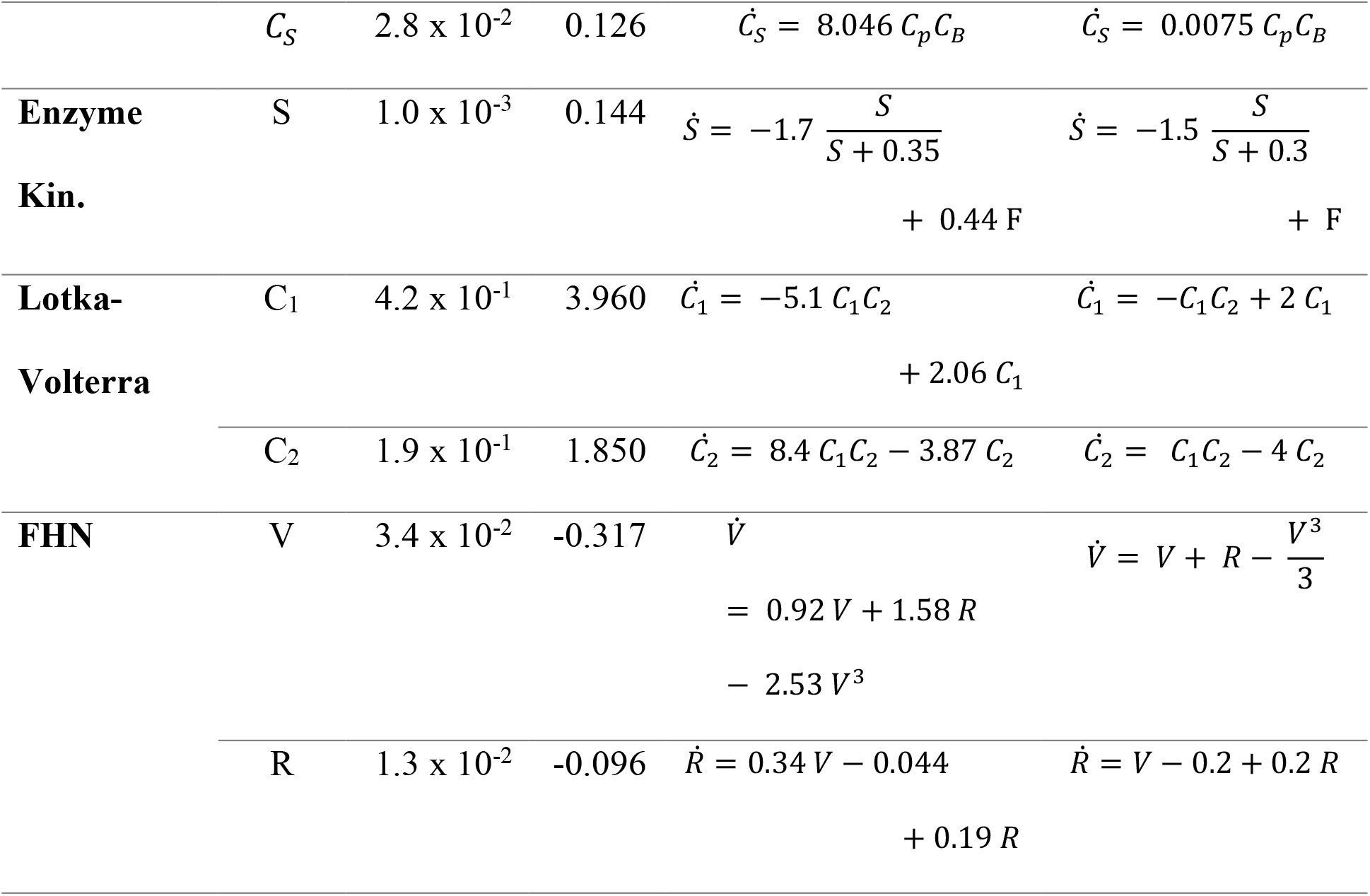
RMSEP made by the Functional-Hybrid model and the final equation deduced by it for the different case studies are compared with the mean statistics of the data and the actual equations used to generate data, respectively.

**Figure 3:**
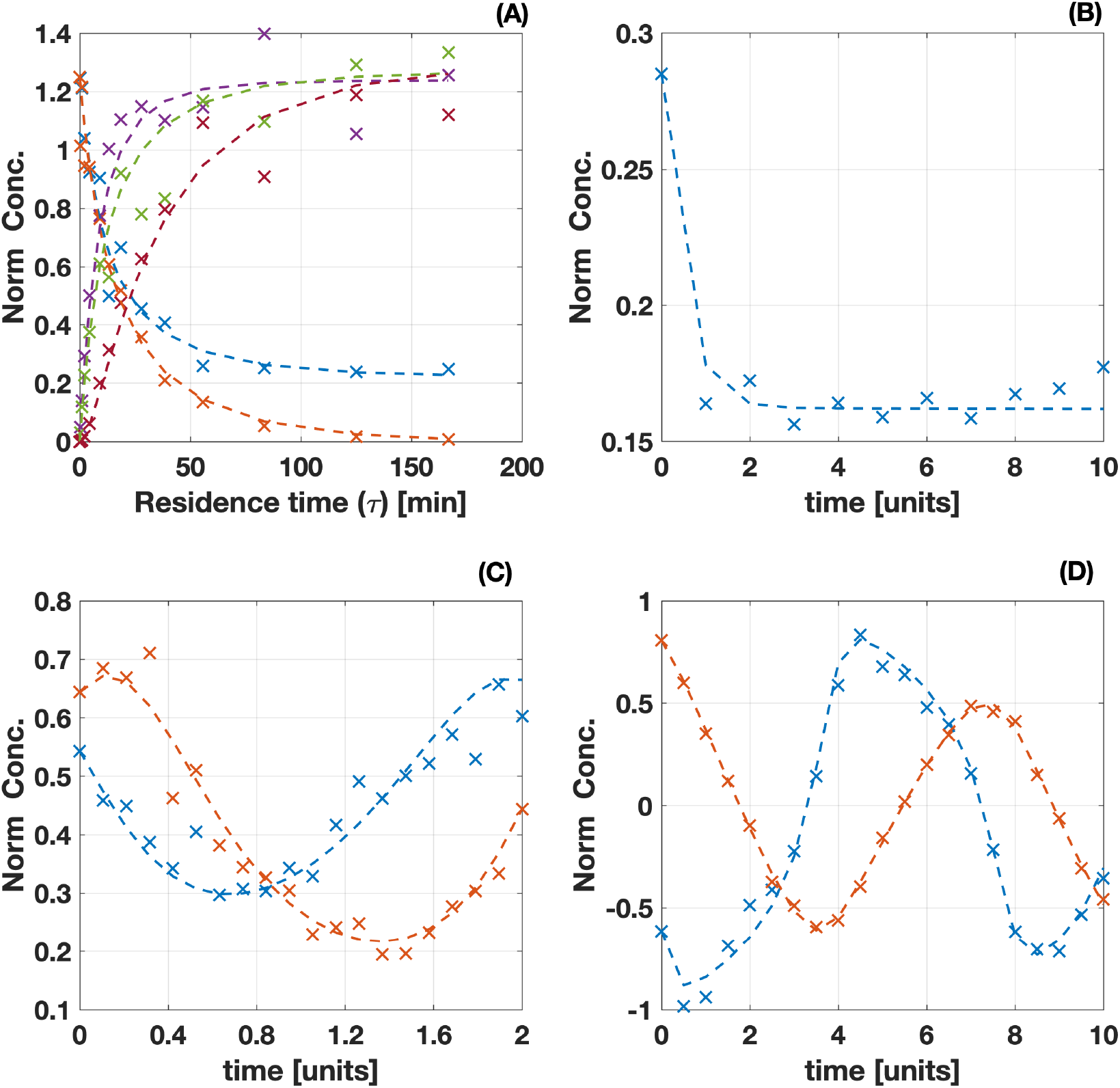
Time evolution of the normalized concentrations (to the maximum value in the training data) of the (A) five chemical species involved in the reaction as a function of the residence time in the reaction kinetic identification (B) substrate concentration in the Michaelis-Menten case study (C) two states in the Lotka-Volterra system and (D) the two states in the FHN system. The crosses represent the measurements and the dashed line represent the model prediction.

### 3.2. Microbial Fermentation Bioreactor

The results from these four case studies validated the performance of the proposed algorithm. Subsequently, the Function-Hybrid framework was applied to a case study of relevance to the biotech industry, a simulation of an industrial microbial fermentation bioreactor. As detailed in section 2.1.5, 30 experiments were generated using the LHS method. 20 randomly chosen experiments were used as the training set and following the protocols detailed in Section 2.2.1, the Functional-Hybrid model was trained. The final trained (or optimized) model was used to predict the test set consisting of the remaining 10 experiments. Figure 4A shows the comparison of the true value (black solid line), the perturbed measurements (blue crosses) and the predictions (red dashed lines) made by the Functional-Hybrid model for all the three states, X, S and P, on an exemplary run in the test set. It is observed that the dynamic profiles predicted by the Functional-Hybrid model overlaps the true simulated profiles for all the three variables despite being trained using the perturbed noisy measurements. The strict structural relationship imposed through domain-specific transformations alleviates the model from being influenced by noise and subsequently overfitting to the noise.

**Figure 4:**
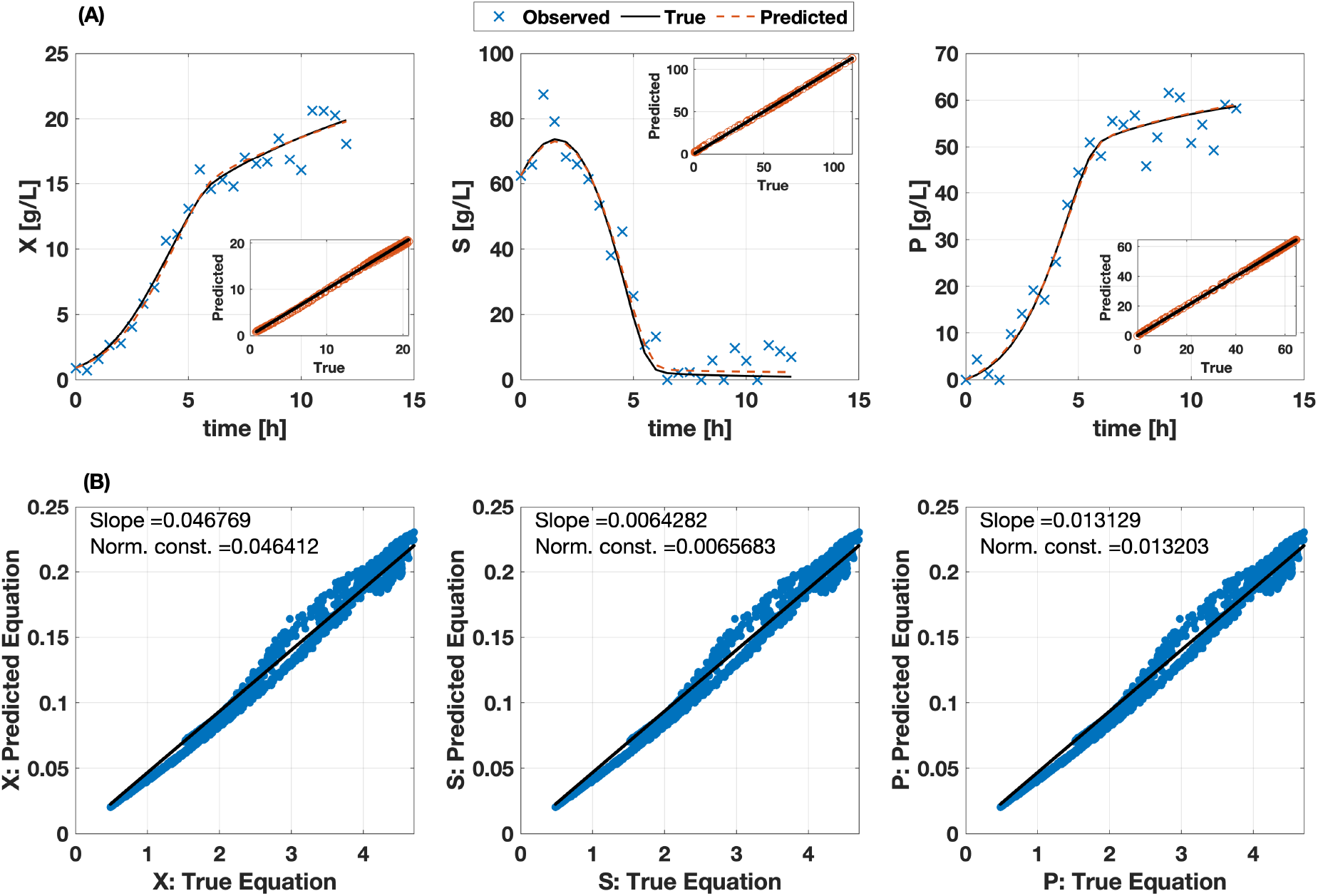
(A) Comparison of the true (black solid), measured (blue cross) and Functional-Hybrid model predicted (red dashed) time profile of an exemplary run from the test set. Parity plots of true values (x-axis) and Functional-Hybrid model predicted values (y-axis) for all the experiments in the test set is presented in bottom-right, top-right and bottom-right corner of the plots for the three states X, S and P, respectively. (B) The parity plot of true equation (x-axis) and Functional-Hybrid model predicted equation (y-axis) for several randomly simulated combinations of X, S and P.

The parity plots presented within the subplots in Figure 4A shows the comparison of true values (not the measured or perturbed values) and the predicted values of the respective states X, S and P, for all the experiments in the test set. The parity plots do not show any systematic bias or any large variation along the diagonal indicating the perfect correlation between the true and predicted values. This confirms that the behavior, i.e., the predictions aligning closely with the true values, depicted by the exemplary dynamic profile is similar in all the other experiments in the test set as well.

In addition to accurate predictions of the dynamic profiles, the system of ODE equations learnt by the Functional-Hybrid model estimates the values for the three equations 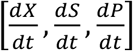 coherent with the values derived from the original in-silico model. Figure 4B demonstrates the correlation of the values of the equation for several randomly simulated combinations of the states [X, S and P] derived from the original model (x-axis) and those estimated by the Functional-Hybrid model (y-axis). Though there are slight deviations among the two, the Functional-Hybrid model was capable of learning the representation of the equations that resembles the original system of equations. These deviations could be attributed to two factors: (i) the Functional-Hybrid model was trained using noisy measurements, which is more realistic as the true system can be observed only with some noise, and (ii) the numerical approximations due to the type of optimizers used.

Additionally, it is here to be noted that the scale of the x and y axis in the plots of Figure 4B are different and that the slope of the correlation line is 4.67 × 10^−2^, 6.42 × 10^−3^ and 1.31 × 10^−2^ for the respective three states instead of 1. This is due to the fact that the Functional-Hybrid model is learnt based on normalized concentrations of the states while the original system of ODE is based on un-normalized concentrations. Thus, as indicates in Figure 4B, the slope of the correlation line is equal to the normalization constant used to scale the respective states. The finally converged Functional-Hybrid model learns the following equations for the three states:

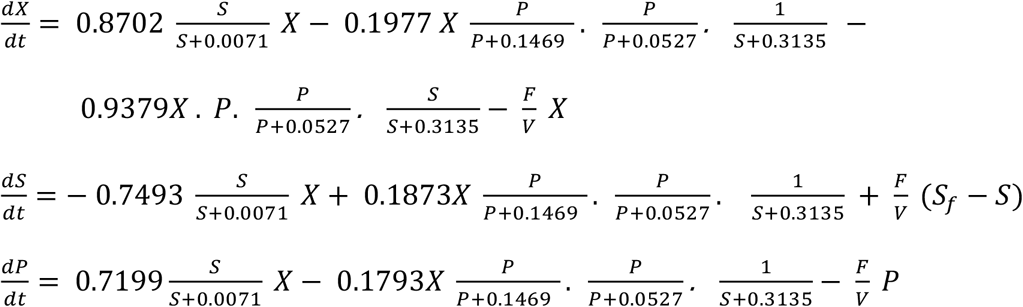

Though the converged equations do not have the exact same functional representation as the original system of ODEs, Figure 4B demonstrates that the value of the equations estimated by the converged Functional-Hybrid model correlate well with the original system of equations. The difference in the functional representation learnt by our modeling framework could be attributed to (i) that it is trained on noisy data, (ii) the use of local optimizer in the Functional-Hybrid framework, (iii) the two functional representations are numerically equivalent, which seems plausible from the Figure 4B, (iv) the set of experiments used for model training cannot distinguish between the two forms and specific experiments might be required to distinguish the system of ODE derived by Functional-Hybrid model from the original system of ODE. In this regard, further model discrimination analysis [43] can be used to increase the reliability on the selected structures by generating data that assures structural identifiability [43].

### 3.3. Comparison with the Hybrid-ANN model

The performance of the Functional-Hybrid model is compared with the benchmark approach of developing hybrid models, by using an ANN, for the microbial fermentation bioreactor case study presented in the previous section. The same 20 experiments considered to build the Functional-Hybrid model are used to train the Hybrid-ANN using the methodology detailed in Section 2.2.2. Subsequently the same test set is used to evaluate the performance of both the models. Figure 5 illustrates the same analysis as in Figure 4 but now for the Hybrid-ANN model.

**Figure 5:**
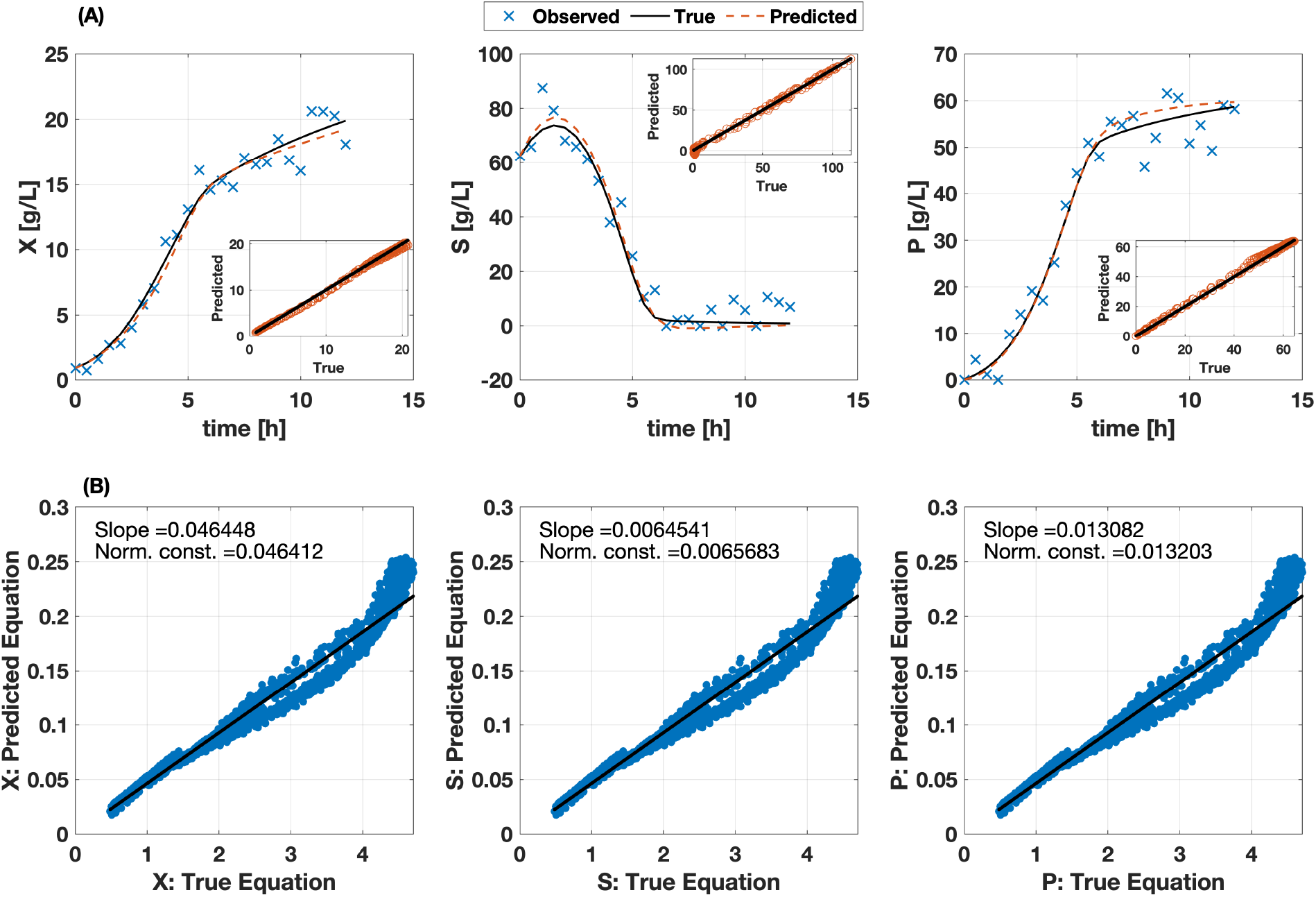
(A) Comparison of the true (black solid), measured (blue cross) and Hybrid-ANN model predicted (red dashed) time profile of an exemplary run from the test set. Parity plots of true values (x-axis) and Hybrid-ANN model predicted values (y-axis) for all the experiments in the test set is presented in bottom-right, top-right and bottom-right corner of the plots for the three states X, S and P, respectively. (B) The parity plot of true equation (x-axis) and Hybrid-ANN model predicted equation (y-axis) for several randomly simulated combinations of X, S and P.

It could be observed in Figure 4A that the Functional-Hybrid model does not show any systematic biases or large variances indicating almost a perfect correlation between the true and predicted values. These predictions thus resulted in very low RMSEP 0.24 g/L, 1.15 g/L and 0.28 g/L for the states X, S and P, respectively. In contrast, the RMSEP attained by the Hybrid-ANN model is 0.45 g/L, 2.25 g/L and 1.35 g/L for X, S and P, respectively which is double that of the Functional-Hybrid model for × and S, and almost five times for the state P. As shown in parity-plots of Figure 5A, the Hybrid-ANN model consistently underpredicts the concentration of × and overpredicts the concentration of P for all the experiments in the test set. While for the state S, the Hybrid-ANN model shows a higher variance throughout the entire concentration range and also predicts unrealistic concentrations, i.e., negative values of concentration. These observations are reflected in higher RMSEPs of the Hybrid-ANN compared to the Functional-Hybrid model for all the three states.

While the equations estimated by Functional-Hybrid model show a very good correlation with the original system of equations (c.f. Figure 4B), the equations deduced by the Hybrid-ANN model show deviations from the numerical values of the original equation (c.f. Figure 5B). At lower concentration, there is good agreement between the deduced and original equation, but with increasing concentration the variance between the predicted and true equation increases considerably. Additionally, at higher concentration there is a systematic bias in the predicted equation which overestimates the true equation. These trends hold true for all the three states:X, S and P.

### 3.4. Practical advantages of Functional-Hybrid model

After performing a base case comparison between the two models, the Functional-Hybrid and the Hybrid-ANN model, the number of experiments required to train the models and their performance in extrapolation was studied. For these studies, again the microbial fermentation bioreactor was considered. As described in Section 2.1.5, 50 experiments were designed using the LHS method and a test set of 10 experiments were randomly chosen among these experiments, and kept fixed.

To study the number of experiments required to train a reliable model, from the remaining 40 experiments, training sets of different sizes were generated using a randomly chosen subset of experiments. The exact same training and test set was used to train the two models. Figure 6A shows the comparison in terms of *MSEP*_*norm,max*_, calculated as per the definition introduced in the Methods section, for the two models trained using different number of experiments. It can be observed that the Functional-Hybrid model requires about 10-20 experiments to reach a very accurate performance while the Hybrid-ANN requires about 40 experiments to reach the same level of performance attained by the Functional-Hybrid model with 20 experiments (highlighted through the zoomed view presented within Figure 6A). The prior domain knowledge encoded in the form of transformations helps the Functional-Hybrid model to converge to a good model with much lesser number of experiments in comparison the Hybrid-ANN that requires more experiments to learn all the dependencies accurately. Since a lot of resource is involved in the experiments and analytics, the reduction in the experimental effort by 50% is of great value to the biotech industry.

**Figure 6:**
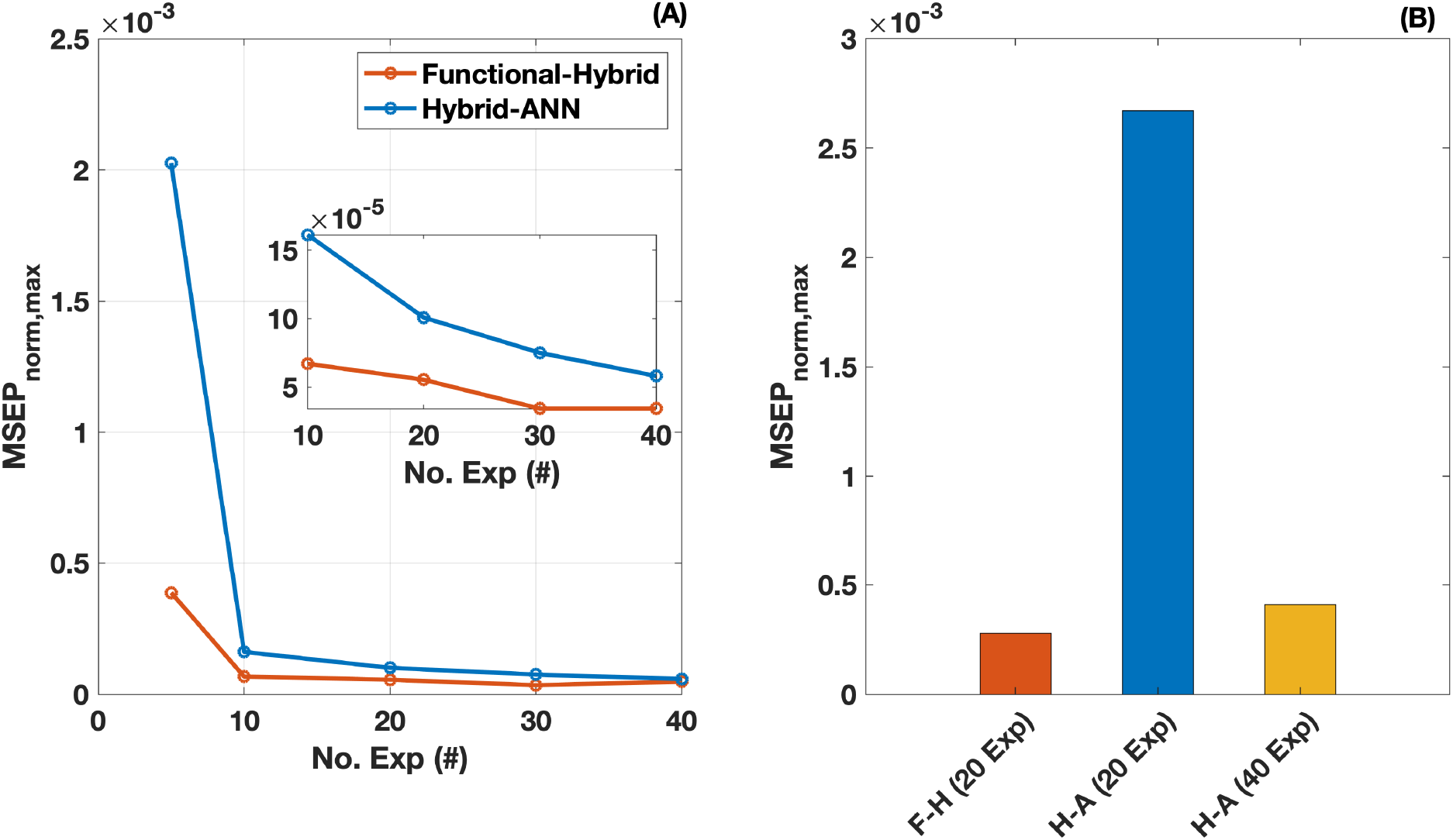
(A) Comparison of the normalized MSEP score of the Functional-Hybrid model and the Hybrid-ANN trained using different number of experiments in interpolation. (B) Comparison of the Functional-Hybrid model trained using 20 experiments (F-H (20 Exp)), Hybrid-ANN trained using the same 20 experiments (H-A (20 Exp)) and Hybrid-ANN trained using 40 experiments (H-A (40 Exp), the point at which the performance of Hybrid-ANN approaches Functional-Hybrid in interpolation, shown in Figure 6 A) based on normalized MSEP score.

Additionally, the strong structures imposed in the Functional-Hybrid model also makes it very robust in its extrapolation capabilities. Figure 6B compares the metric *MSEP*_*norm,max*_ for extrapolation experiments made by the different models namely, Functional-Hybrid model trained with 20 experiments, Hybrid-ANN model trained with 20 experiments and a Hybrid-ANN model trained with 40 experiments (i.e., the number of experiments where the performance of the Hybrid-ANN matches the Functional-Hybrid model in the case of interpolation).

The Hybrid-ANN trained with 20 experiments perform very poor in extrapolation with the *MSEP*_*norm,max*_ being an order of magnitude higher than the ones observed with the Functional-Hybrid model. The *MSEP*_*norm,max*_ of the Functional-Hybrid and Hybrid-ANN model, both trained with 20 experiments, in interpolation is 5.53 × 10^−5^ and 1 × 10^−4^, respectively, resulting in a difference of only 4.54 × 10^−5^. However, the *MSEP*_*norm,max*_ of the Functional-Hybrid and Hybrid-ANN models when extrapolating is 2.78 × 10^−4^ and 0.0027, respectively, resulting in a difference of an order of magnitude. This indicates that the ability to learn the true trend in the training can significantly worsen during extrapolation, thereby leading to poor performance and meaningless results if the models are used for process optimization. In order for such models to be useful for extrapolation, they have to reach sufficiently good training performances, thus, requiring at least 40 experiments during the training. Despite that, the Hybrid-ANN model shows a *MSEP*_*norm,max*_ of 4.1 × 10^−4^, which is about 1.5 times higher in comparison to that of the Functional-Hybrid model. On the other hand, the Functional-Hybrid model produces the most accurate estimates while using the lesser number of experiments, making this model suitable for process optimization and also for its integration with real-time measurements to facilitate monitoring and control [44].

## 4. Conclusion

Hybrid models that are capable of integrating engineering know-how (first-principle models) with data (machine learning) have become a pragmatic solution to modeling in different areas of chemical engineering and biotechnology. However, the data-driven part of these hybrid models is largely dependent on conventional machine learning methods, such as artificial neural network, support vector machines, gaussian processes etc., thus making it difficult for process engineers to interpret the patterns learnt by these models. On the other hand, there are common functional forms specific to each domain that are easily recognized by the experts.

This work presented a novel hybrid modeling framework, Functional-Hybrid models, that uses an adapted symbolic regression strategy based on domain-specific ranked functional forms to build dynamic models that are easier to interpret by domain experts. The framework is successfully implemented for four benchmark systems and a system of relevance to process engineers, a microbial fermentation bioreactor. The developed Functional-Hybrid model is compared against a conventional hybrid model based on artificial neural network (Hybrid-ANN). The models are compared based on its accuracy in interpolation and extrapolation where we could demonstrate that the error of the Functional-Hybrid model is at least 2-times lower than in the Hybrid-ANN. Further, the experimental burden of developing these models was evaluated in terms of number of experiments required to build a robust model for either case, with Functional-Hybrid models requiring only 20 experiments and Hybrid-ANN needing at least 40 experiments. The additional structure enforced by the domain specific functional transformations in the Functional-Hybrid model enhances the robustness of the predictions, specifically when extrapolating, while reducing the experimental data needed to train such models.

To demonstrate the concept, the case studies presented in this work are based on simulations. However, addressing experimental cases (not based on in silico data) are the foreseen next steps. Additionally, the current framework could be improved in terms of computational performance, which constitutes another future research direction.

## Supplementary Information

System of equations used to simulate data for the different case studies.

### 1. Chemical reaction system

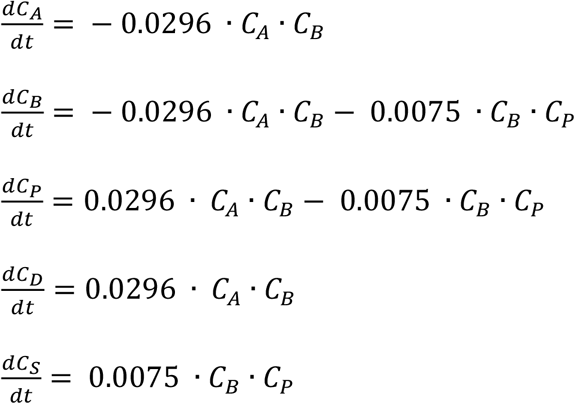

where *C*_*A*_, *C*_*B*_, *C*_*P*_, *C*_*D*_ and *C*_*S*_ are the concentration of different chemical species A, B, P, D and S.

**S.I. Table 1:** Ranked transformations used as input to the algorithm, stepwise addition iteration where optimized model meets threshold (Converged) and the optimal number of blocks (*N*_*Blocks*_) for the chemical reaction kinetics case study.

| Rank | Transformation | Entity | Converged | $N_{Blocks}$ |
| --- | --- | --- | --- | --- |
| 1. | Identity | $C_A, C_B, C_P, C_D, C_S$ | Yes | 2 |
| 2. | Power law | $(C_A)^{k_1}, (C_B)^{k_2}, (C_P)^{k_3}, (C_D)^{k_4},$<br>$(C_S)^{k_5}$ | - | - |
| 3. | Monod | $\frac{C_A}{C_A+k_6}, \frac{C_B}{C_B+k_7}, \frac{C_P}{C_P+k_8}, \frac{C_D}{C_D+k_6}, \frac{C_S}{C_S+k_6}$ | - | - |

### 2. Enzyme Kinetics

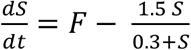

where S is the substrate concentration and F is the inflow flux of the substrate.

**S.I. Table 2:**
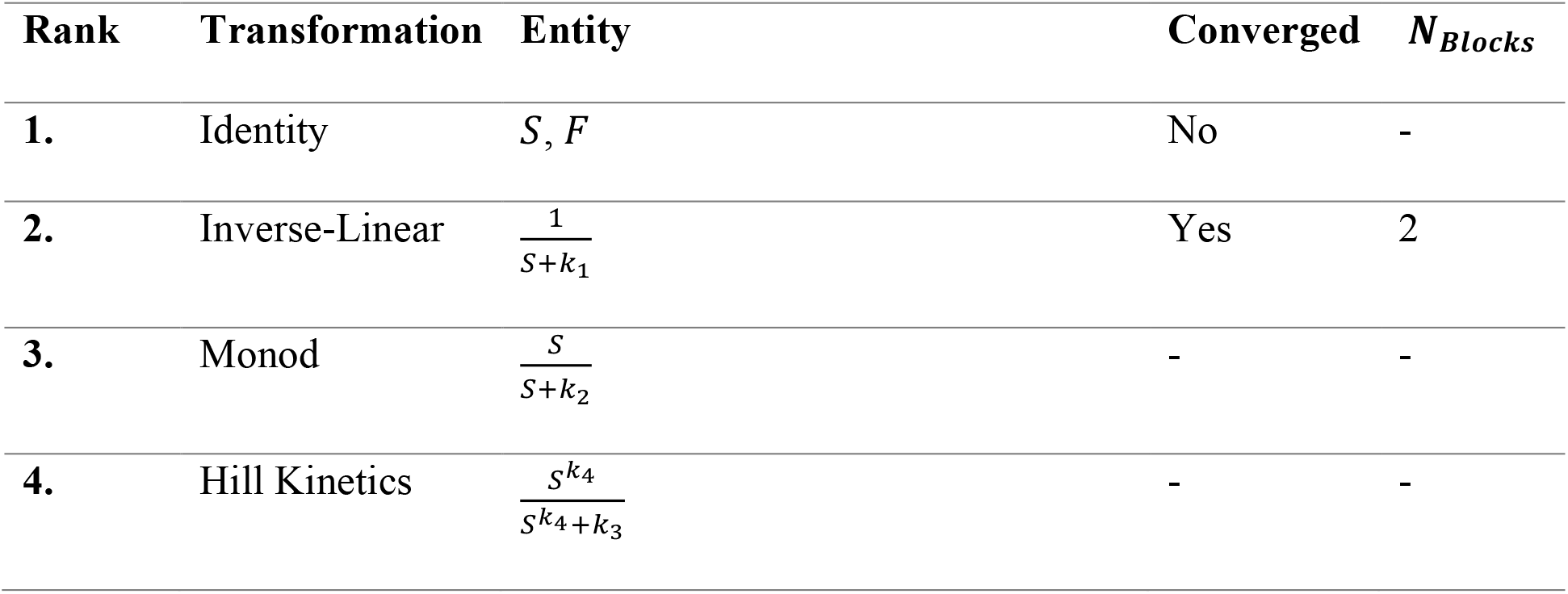
Ranked transformations used as input to the algorithm, stepwise addition iteration where optimized model meets threshold (Converged) and the optimal number of blocks (*N*_*Blocks*_) for the enzyme kinetics case study.

### 3. Lotka-Volterra System

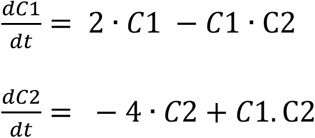

where C1 and C2 are the concentrations of the two states involved in the Lotka-Volterra system.

**S.I. Table 3:** Ranked transformations used as input to the algorithm, stepwise addition iteration where optimized model meets threshold (Converged) and the optimal number of blocks (*N*_*Blocks*_) for the Lotka-Volterra problem.

| Rank | Transformation | Entity | Converged | $N_{Blocks}$ |
| --- | --- | --- | --- | --- |
| 1. | Identity | $C_1, C_2$ | Yes | 3 |
| 2. | Quadratic | $(C_1)^2, (C_2)^2$ | - | - |
| 3. | Sigmoid | $\frac{C_1}{C_1+k_1}, \frac{C_2}{C_2+k_2}$ | - | - |

### 4. FitzHugh-Nagumo (FHN) System

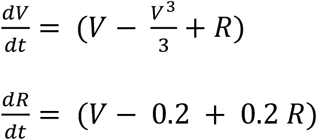

where V and R are the concentrations of the two states involved in the FHN system.

**S.I. Table 4:**
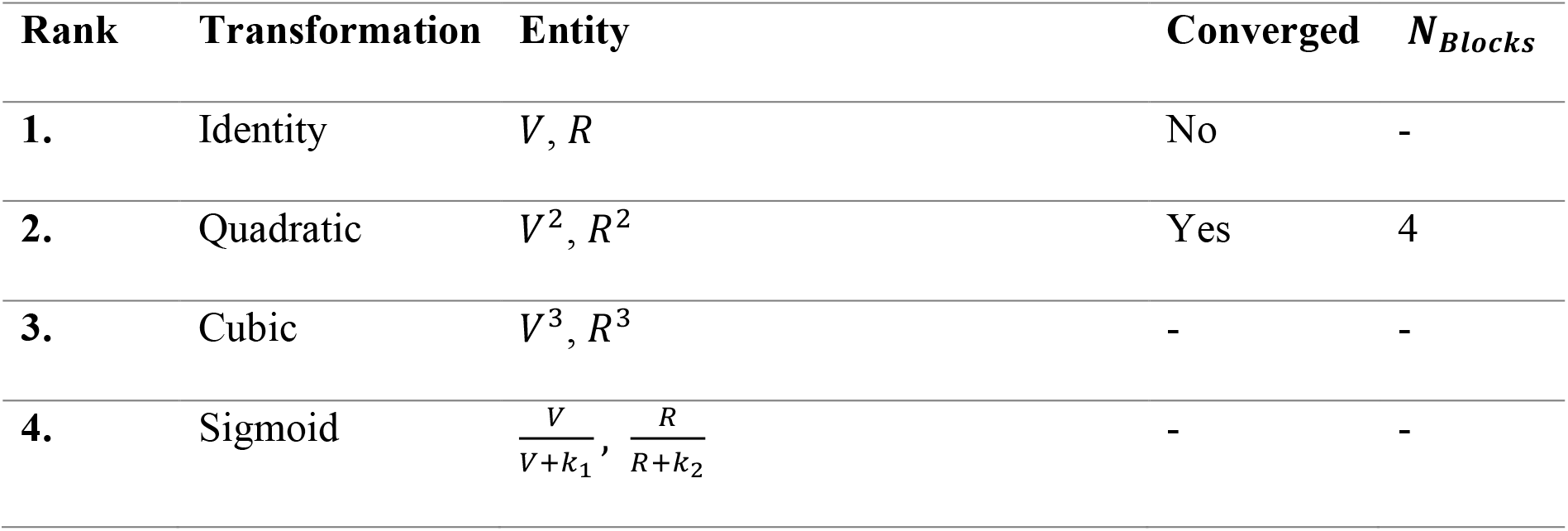
Ranked transformations used as input to the algorithm, stepwise addition iteration where optimized model meets threshold (Converged) and the optimal number of blocks (*N*_*Blocks*_) for the FHN system.

### 5. Microbial Fermentation Bioreactor

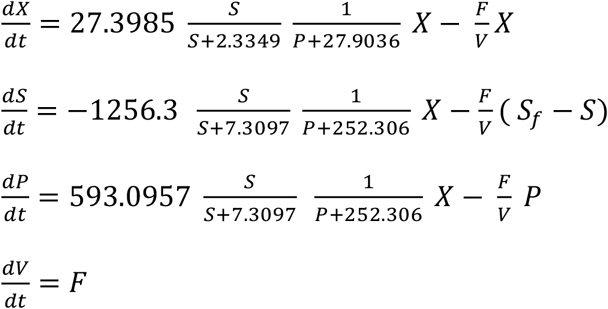

where X, S and P are the concentration of biomass, substrate and product, F is the feed rate and *S*_*f*_ is substrate concentration of the feed.

**S.I. Table 5:** Ranked transformations used as input to the algorithm, stepwise addition iteration where optimized model meets threshold (Converged) and the optimal number of blocks (*N*_*Blocks*_) for the Microbial Fermentation case study.

| Rank | Transformation | Entity | Converged | $N_{Blocks}$ |
| --- | --- | --- | --- | --- |
| 1. | Identity | $X, S, P$ | No | - |
| 2. | Sigmoid | $\frac{S}{S+k_1}, \frac{P}{P+k_2}$ | No | - |
| 3. | Inverse-Linear | $\frac{1}{S+k_3}, \frac{1}{P+k_4}$ | Yes | 3 |
| 4. | Inverse-Linear | $\frac{1}{S+k_5}, \frac{1}{P+k_6}$ | - | - |
| 5. | Inverse-Quadratic | $\frac{1}{S+k_7+k_8S^2}, \frac{1}{P+k_9+k_{10}P^2}$ | - | - |

**S.I. Table 6:** RMSEP made by the Functional-Hybrid model for the different case study compared with the min, max and mean statistics of the data and the final equations deducted by the Functional-Hybrid model for the different cases.

| Case | States | RMSEP<br>[units] | Min<br>[units] | Max<br>[units] | Mean<br>[units] | Equation |
| --- | --- | --- | --- | --- | --- | --- |
| Reaction Kin. | $C_A$ | 0.0054 | 0.38 | 2 | 1.145 | $\dot{C}_A = -9.6 C_A C_B$ |
| | $C_B$ | 0.0092 | 0.03 | 2 | 1.015 | $\dot{C}_B = -1.38 C_P C_B$<br>$- 9.65 C_A C_B$ |
| | $C_P$ | 0.0150 | 0.0 | 1.24 | 0.723 | $\dot{C}_P = -2.62 C_P C_B$<br>$+ 15.6 C_A C_B$ |
| | $C_D$ | 0.0067 | 0.0 | 1.62 | 0.864 | $\dot{C}_D = 11.93 C_A C_B$ |
| | $C_S$ | 0.0282 | 0.0 | 0.36 | 0.126 | $\dot{C}_S = 8.046 C_P C_B$ |
| **Enzyme Kin.** | S | 0.001 | 0.0252 | 0.939 | 0.1443 | $\dot{S} = -1.7 \frac{S}{S + 0.35} + 0.44 F$ |
| **Lotka-** | C<sub>1</sub> | 0.42 | 1.36 | 8.76 | 3.96 | $\dot{C}_1 = -5.1 C_1 C_2 + 2.06 C_1$ |
| **Volterra** | C<sub>2</sub> | 0.197 | 0.38 | 5.00 | 1.85 | $\dot{C}_2 = 8.4 C_1 C_2 - 3.87 C_2$ |
| **FHN** | V | 0.0342 | -2.823 | 2.909 | -0.317 | $\dot{V} = 0.92 V - 1.58 R$<br>$- 2.53 V^3$ |
| | R | 0.013 | -3.221 | 4.607 | -0.096 | $\dot{R} = 0.34 V - 0.044 + 0.19 R$ |

